# *De novo* protein discovery in non-model organisms

**DOI:** 10.64898/2026.05.08.723910

**Authors:** Asif Ali

## Abstract

We developed *plant* (Parallel Annotation of Transcriptomes), a *de novo* method that can potentially compare RNA-seq data of any two species without a reference genome. *plant* is conceptually similar to chromatography. In the same way a complex mixture is filtered to isolate its individual components, we applied a computational method to identify, annotate, and quantify components across transcriptomes. The comparison points are universal protein domain annotations rather than species-specific genes, as would be the case for a differential gene expression analysis. We looked at several *Selaginella* species via the 1000 Plant transcriptomes initiative (1KP) where RNA-seq data for various plant species have been made publicly available. The raw reads were assembled via Trinity. The assembled transcripts were then searched against the Pfam protein domain database via InterProScan. The assembled transcripts were also quantified via kallisto. By merging these two aspects, we were able to see how often a particular protein domain – a predicted protein structure – is expressed. These quantified annotations of protein domains are comparable across species, assuming a relatively short evolutionary distance. We were also able to identify the presence of species-specific protein domains and trace each annotation back to the gene. A bubble plot was created to visualize the distributions of Pfam annotations across species as well as GO terms.

## Introduction

Plants occupy a wide variety of ecological roles across the biosphere today. All green plants are capable of photosynthesis, a process by which atmospheric carbon is captured and sugar is synthesized via sunlight. Photosynthesis takes place in the chloroplast, a double membranous organelle that was once an engulfed cyanobacteria. This endosymbiosis was maintained over evolutionary time through vertical transmission, creating an environment for gene exchange and protein trafficking whereby the cyanobacteria eventually became incorporated as a crucial component to form a single unit – the first plant cell^1,2^.

Green plants, which include green algae and terrestrial plants, have the same coupled pigments, chlorophyll *a* and *b*, to capture light for carbon sequestration^3^. All terrestrial plants evolved from an aquatic environment. Green algae, charophytes in particular, resemble a link to bryophytes, a clade where features of early terrestrial plants are conserved^2^. Angiosperms, or flowering plants, occupy the top position along the phylogenetic tree where we see the most derived features (Figure 1). The most studied plant in modern science, *Arabidopsis thaliana*, flowers^4^. There are over 200 other flowering plants with sequenced genomes^5^. Angiosperms, gymnosperms, and ferns all possess megaphylls, or true leaves, and form the euphyllophytes. Lycophytes have microphylls and occupy a key phylogenetic position as sister to all other vascular plants (Figure 1). Lycophytes include clubmosses as well as the genera *Isoetes* and *Selaginella* – the quillworts and spikemosses, respectively^6^.

**Figure 1.**
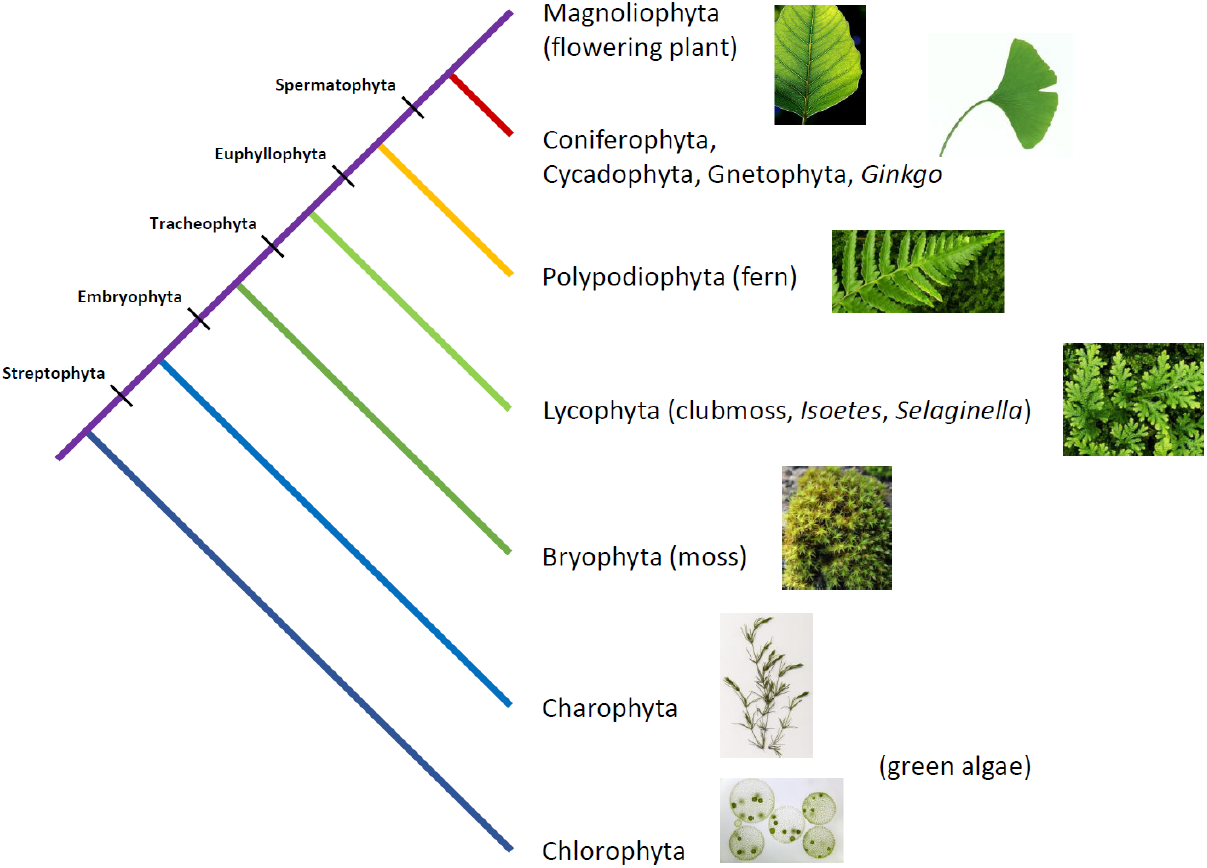
Phylogeny of Viridiplantae as part of the Open Tree of Life^46^. Updated. Charales image by Fischer^47^.

The genome of a lycophyte, *Selaginella moellendorffii*, has been sequenced^6^, revealing genus-specific genes associated with alternation of generations^7^. Subsequent studies on RNA-seq data have found differential expression among tissue types^8^. RNA-seq data of a diverse array of green plants, including other *Selaginella* species, have recently been made publicly available by the 1000 Plant transcriptomes initiative (1KP). Collaborators assembled the reads *de novo* and include associated protein-protein interactions mapped to metabolic pathways through an interactive webpage^9^. Additionally, extensive work has been done to assess the phylogeny of shared gene families across the green plant lineages^10^.

We developed a novel application of existing methods – *plant* (Parallel Annotation of Transcriptomes). First, we independently assemble the reads *de novo*. We then annotate the assembled reads. We also pseudoalign the raw reads to the assembled reads to quantify the assembled reads. By merging these two aspects, we are able to quantify the annotations to use as universal comparison points rather than genes. This approach allows us to compare closely related species without a reference genome.

Annotations that are unique to a species may indicate novel protein discovery. We include an interactive visualization to identify species-specific annotations.

## Samples

The *Selaginella* species sampled by 1KP collaborators include *S. lepidophylla, S. wallacei, S. willdenowii, S. selaginoides, S. apoda, S. kraussiana, S. acanthonota*, and *S. stauntoniana*^9^. Native to the Chihuahuan Desert of Northern Mexico, New Mexico, and Texas as well as Baja California, Southern Mexico, El Salvador, and Costa Rica, *S. lepidophylla* is a medicinal herb and a “resurrection plant” capable of poikilohydry^11,12,13^. *S. wallacei* is native to Alberta, British Columbia, California, Idaho, Montana, Oregon, and Washington. Native to Cambodia, Indonesia, Laos, Malaysia, Myanmar, the Philippines, Thailand, and Vietnam, *S. willdenowii* is adapted to shade with a blue iridescence to capture light beneath the canopy^14^. Present across the Northern Hemisphere, *S. selaginoides* is a circumboreal species sister to all other sampled *Selaginella* species^15^ (Figure 2). *S. apoda* is native to Eastern United States, Northeastern Mexico, along the Golf of Mexico, Southern Mexico, Guatemala, and Cuba^16^. *S. kraussiana* is native to the Tsitsikamma Forests of the Eastern Cape, KwaZulu-Natal, Mpumalanga, and Limpopo, tropical regions of Eswatini, Botswana, Zimbabwe, Zambia, Mozambique, Malawi, Tanzania, Burundi, Rwanda, Uganda, Kenya, Ethiopia, Sudan, Angola, Zaïre, Congo, Cameroon, Equatorial Guinea, and Sierra Leone, as well as the islands of Bioko, Comoros, Madagascar, Mauritius, the Azores, Madeira, and the Canary Islands^17^. *S. acanthonota* is native to North and South Carolina, Florida, and Georgia. *S. stauntoniana* is native to China, Korea, Mongolia, and Taiwan^18^.

**Figure 2.**
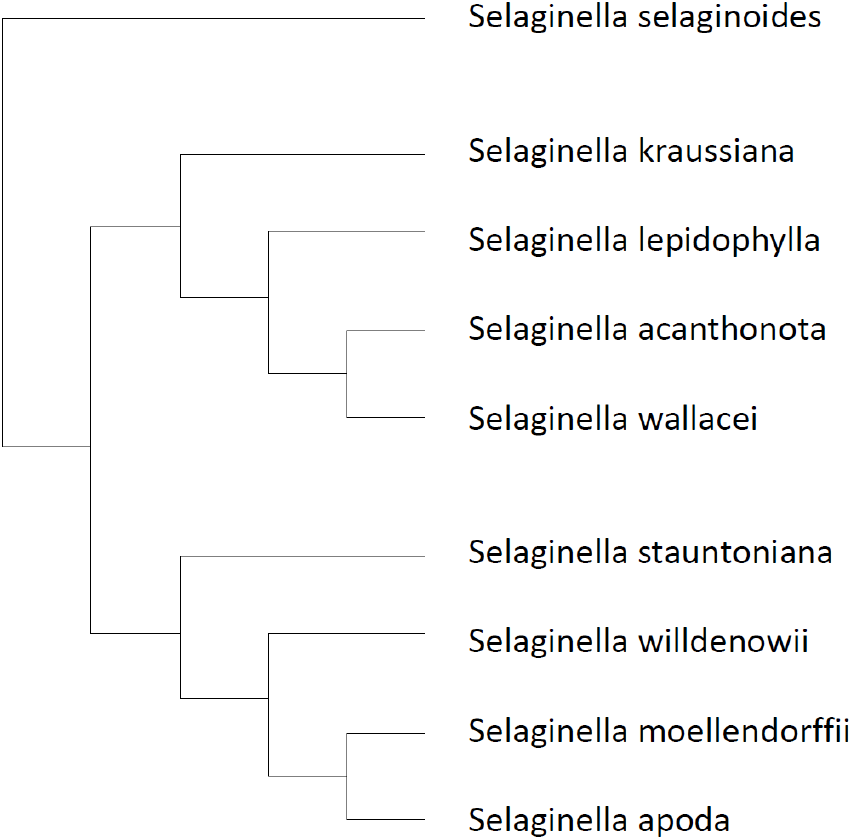
Phylogeny of the *Selaginella* species sampled by 1KP collaborators. Information was derived from the phylogenetic tree in the American Journal of Botany. The paper – “Phylogeny of Selaginellaceae: there is value in morphology after all!” – linked morphological adaptations among various *Selaginella* species. The authors looked at chloroplast and nuclear gene sequence to construct a detailed phylogenetic tree^15^.

The plant specimens were sampled for RNA-seq data. All samples appear to be purely vegetative tissue. The *S. wallacei* sample (JKAA), however, may include reproductive tissue as the authors note a whole “leaf” structure. Additionally, we should note that *S. stauntoniana* (ZZOL) had its gemmae sampled along with both vegetative and reproductive tissue^19^. A gemma is a clone of the parent plant formed through asexual propagation (Table 1). The raw RNA-seq reads are available through the Sequence Read Archive (SRA)^20^. OneKP collaborators have already assembled the raw RNA-seq reads via SOAPdenovo-Trans^9^. We independently assemble the raw RNA-seq reads via Trinity. We then annotate and quantify the transcripts via InterProScan and kallisto, respectively. Finally, we merge the quantitative information with the annotations.

**Table 1.** A list of sample information for the *Selaginella* species selected by 1KP collaborators^9^.

| OneKP identification | Tissue of RNA-seq sample | Species of *Selaginella* |
| --- | --- | --- |
| ABIJ | young fronds | *S. lepidophylla* |
| JKAA | leaf | *S. wallacei* |
| KJYC | tips of shoots | *S. willdenowii* |
| KUXM | tips of shoots | *S. selaginoides* |
| LGDQ | sterile leaves/branches | *S. apoda* |
| ZFGK | tips of shoots | *S. kraussiana* |
| ZYCD | sterile leaves/branches | *S. acanthonota* |
| ZZOL | sterile and fertile material, and gemmae | *S. stauntoniana* |

## Methods

We developed *plant* (Parallel Annotation of Transcriptomes) to process the data. *plant* is a bioinformatics pipeline that is conceptually similar to a differential gene expression analysis; the difference is that the distributions of each annotation are quantified and compared rather than species-specific genes. Annotations are universally comparable, whether GO terms or Pfam protein domains. *plant* also has the potential to indirectly discover new proteins by locating annotations that are unique to a transcriptome or species. *plant* is comprised of a series of open-source bioinformatics software. We assembled the RNA-seq reads as well as annotated the assembled transcripts. We then quantified the annotations by quantifying each transcript independently – each transcript was quantified by mapping the raw reads to the assembled scaffolds – and then passing that information down to the annotation level. This abstract form of differential expression – distributions of protein domains across genes and isoforms rather than counts of genes – provides us with a quantification of the annotation per transcript and thus allows us to assess transcript abundance of a potentially undiscovered protein. For protein discovery, the annotation along with an E-value supports the notion that the novel protein domain is actually present. The quantification then ensures that the novel protein domain is predicted to be expressed at a biologically relevant scale.

Initial data processing was done in a Unix environment. With the SRA Toolkit, we downloaded the raw RNA-seq reads for each species as two FASTQ files as the samples were created with paired-end sequencing. FastQC^21^ was then loaded to check the quality of the reads and search for adapter sequences. The raw RNA-seq reads were then assembled *de novo* via Trinity with minimum contig length of 200bp across all species. Trinity is able to reconstruct whole transcriptomes through three steps. First, *Inchworm* creates the main transcripts while noting alternative splicing. Then, *Chrysalis* clusters the various isoforms as genes and creates a Bruijn graph for each gene. Finally, *Butterfly* processes the graphs to assesses alternative splicing as well as gene paralogs^22,23^. The Trinity jobs were sent to the High Performance Computing cluster Prince at New York University.

Next, we ran InterProScan on our Trinity assembled scaffolds as well as their SOAPdenovo-Trans assembled scaffolds. InterProScan is able to predict protein function via sequence annotation^24^. The scaffolds, or transcripts, were virtually translated and searched against the Pfam database of protein families where there are currently just under 18,000 known structures^25^. Each structure is modeled by a multiple sequence alignment generated through a profile hidden Markov model^26,27^ created by an initial seed alignment^28^ and improved by incorporating UniProtKB sequences^29^ of reference proteomes^30^. We also searched for Gene Ontology (GO) terms^31^, another option of InterProScan^24^. Data was split for parallel processing. Prince cluster at New York University took around 7 hours with 320 cores to process assembled scaffolds. InterProScan created a TSV file listing Pfam protein domains associated with each transcript along with an E-value for each prediction. GO terms were also included.

With the annotations complete, we ran kallisto to quantify the transcripts. Here, the raw reads were pseudoaligned with the transcripts. This approach is efficient and could run on a local machine as pseudoalignment preserves abundance information without the need for an actual alignment^32^. We launched kallisto through Trinity. The kallisto jobs were sent to the Prince cluster and produced a TSV file of transcript isoform abundances.

On a local machine in R^33,34^, Pfam protein domain annotations and associated E-values, GO terms, and TPM values were extracted. All data was merged with the help of plyr^35^ and then plotted via ggplot2^36^ and plotly^37^. We link the species-specific GO terms and protein domains with their associated Trinity assembled transcripts. We also filter the protein domain predictions by E-value.

The Unix environment and computational jobs sent to the Prince cluster had resources allocated – particularly RAM – greatly exceeding the amount necessary to run the task. Additional capacity is crucial for big data. The InterProScan runs were CPU-optimized for parallel processing. Protocol available at https://github.com/asifali-bio/plant. *plant* is open-source.

## Results

No adapter sequences were found by FastQC. Reads were very high quality. There was 45 to 55 percent GC content in the paired-end RNA-seq reads across all species (Table 2). For sample ABIJ, the median quality score was above 28 across all read positions. Average quality score only dipped below 28 on the final read position. There appears to be heavy sequence duplication (<70%), perhaps due to PCR amplification cycles or initial sample material. There were several sequences with a disproportionate abundance. For sample JKAA, both the average and median quality score was above 28 across all read positions. The GC distribution curve began with a slight initial peak followed by a much sharper peak than the expected normal distribution. There appears to be moderate sequence duplication (>30%).

**Table 2.** Quality and initial assessment of raw RNA-seq reads.

| OneKP ID | Total reads (raw) | Paired reads (raw) | Poor quality sequences | %GC |
| --- | --- | --- | --- | --- |
| ABIJ | 28267020 | 14133510 | 0 | 54 |
| JKAA | 23843130 | 11921565 | 0 | 50 |
| KJYC | 26315654 | 13157827 | 0 | 52 |
| KUXM | 23564384 | 11782192 | 0 | 45 |
| LGDQ | 27907726 | 13953863 | 0 | 55 |
| ZFGK | 24706378 | 12353189 | 0 | 53 |
| ZYCD | 23444448 | 11722224 | 0 | 54 |
| ZZOL | 29882404 | 14941202 | 0 | 52 |

Repeat A and repeat T were disproportionately abundant. For sample KJYC, both the average and median quality score was above 28 across almost all read positions. Only the final read position had an average quality score at 28. There appears to be heavy sequence duplication (>60%) as well as sequences with a disproportionate abundance. For sample KUXM, both the average and median quality score was above 28 across all read positions. There appears to be heavy sequence duplication (<70%) as well as sequences with a disproportionate abundance. For sample LGDQ, both the average and median quality score was above 28 across all read positions. There appears to be heavy sequence duplication (>60%) as well as one sequence in particular with a disproportionate abundance. For sample ZFGK, both the average and median quality score was above 28 across almost all read positions. Only on the final read position was the average quality score just below 28. There appears to be heavy sequence duplication (>60%) as well as one sequence in particular with a disproportionate abundance. For sample ZYCD, both the average and median quality score was above 28 across all read positions. There appears to be heavy sequence duplication (<60%) as well as sequences with a disproportionate abundance. For sample ZZOL, both the average and median quality score was above 28 across all read positions. There appears to be moderate sequence duplication (<50%). From around 1 to 1.5 million read pairs, we were able to construct around 100,000 assembled scaffolds for most species via Trinity. OneKP collaborators generated almost 50,000 scaffolds per species via SOAPdenovo-Trans. JKAA appears to be a distinct case where nearly 200,000 scaffolds were constructed with both Trinity and SOAPdenovo-Trans (Table 3).

**Table 3.** Assembled reads are the scaffolds. Modern RNA-seq technology collects data on the relative abundance of each transcript present, so all samples properly have an equal total transcript abundance after TPM normalization. Derived protein domain abundances and GO term abundances are per million transcripts and vary, however. All abundances are of Trinity assembled reads.

| OneKP ID | Assembled reads |  | Total transcript abundance – kallisto (TPM) | Derived abundance (per million transcripts) |  |
| --- | --- | --- | --- | --- | --- |
|  | OneKP | Trinity |  | Pfam protein domains | GO terms |
| ABIJ | 40120 | 95111 | 1000000 | 785005.2 | 955886.8 |
| JKAA | 182120 | 194556 | 1000000 | 678887.9 | 783416.2 |
| KJYC | 49842 | 101267 | 1000000 | 804232.9 | 996530 |
| KUXM | 44322 | 102371 | 1000000 | 501623.5 | 544076.7 |
| LGDQ | 44749 | 103062 | 1000000 | 710350.4 | 893949.4 |
| ZFGK | 43036 | 89360 | 1000000 | 819126.1 | 1016133 |
| ZYCD | - | 95644 | 1000000 | 828756.9 | 918268.2 |
| ZZOL | 54465 | 115446 | 1000000 | 1136997.7 | 1161569.1 |

Of the total 4892 unique GO terms among all sampled *Selaginella* species and 5595 unique protein domains, 1111 GO terms and 3469 protein domains are shared among all species. The JKAA sample had a noticeably high number of scaffolds constructed via Trinity and SOAPdenovo-Trans as well as the greatest number of unique GO terms (1397) and unique protein domains present (5050). The ZZOL sample had 1240 unique GO terms and 4310 unique protein domains present, in comparison. The JKAA sample also had by far the greatest number of species-specific protein domains present (655) and GO terms (130), perhaps due to sampled reproductive tissue. The ZZOL sample only had 97 species-specific protein domains present (Table 4). However, the ZZOL sample had the greatest density, or relative abundance, of projected protein domains per million transcripts. To assess absolute abundance, a spike-in transcript would be necessary. If we assume that all transcripts are annotated equally, then we can hypothesize that ZZOL had the greatest total abundance of transcripts, perhaps due to sampled reproductive tissue and gemmae, which may have progenitor and stem cell activity as well as various proteins allocated to organ differentiation while under active propagation. If we further assume that all transcripts are translated equally, then we can also hypothesize that ZZOL had the greatest total abundance of translated proteins. The KUXM sample, on the other hand, had the lowest density, or relative abundance, of projected protein domains per million transcripts (Table 4).

**Table 4.** Total count of unique protein domains and GO terms for each species. Also included are species-specific protein domains and GO terms as well as their respective genes.

| OneKP ID | Total unique protein domains (Pfam) | Species-specific |  | Total unique Gene Ontology (GO) | Species-specific |  |
| --- | --- | --- | --- | --- | --- | --- |
|  |  | Protein domains (Pfam) | Genes |  | GO terms | Genes |
| ABIJ | 4076 | 28 | 38 | 1207 | 3 | 3 |
| JKAA | 5050 | 655 | 1976 | 1397 | 130 | 291 |
| KJYC | 4120 | 37 | 52 | 1214 | 6 | 8 |
| KUXM | 4007 | 39 | 75 | 1203 | 9 | 15 |
| LGDQ | 3974 | 32 | 45 | 1190 | 0 | 0 |
| ZFGK | 4175 | 50 | 84 | 1238 | 16 | 24 |
| ZYCD | 4241 | 34 | 63 | 1221 | 4 | 6 |
| ZZOL | 4310 | 97 | 128 | 1240 | 9 | 14 |

GO terms and Pfam protein domains of the *de novo* assembled reads were fetched, quantified, and plotted. The visualization includes quantitative information about the distribution of functional protein domains predicted to be present in various *Selaginella* species (Figures 3 and 4). Each species appears to have a unique signature of predicted protein domains across all actively expressed genes. We note all species-specific GO terms and protein domain annotations associated with each gene.

**Figure 3.**
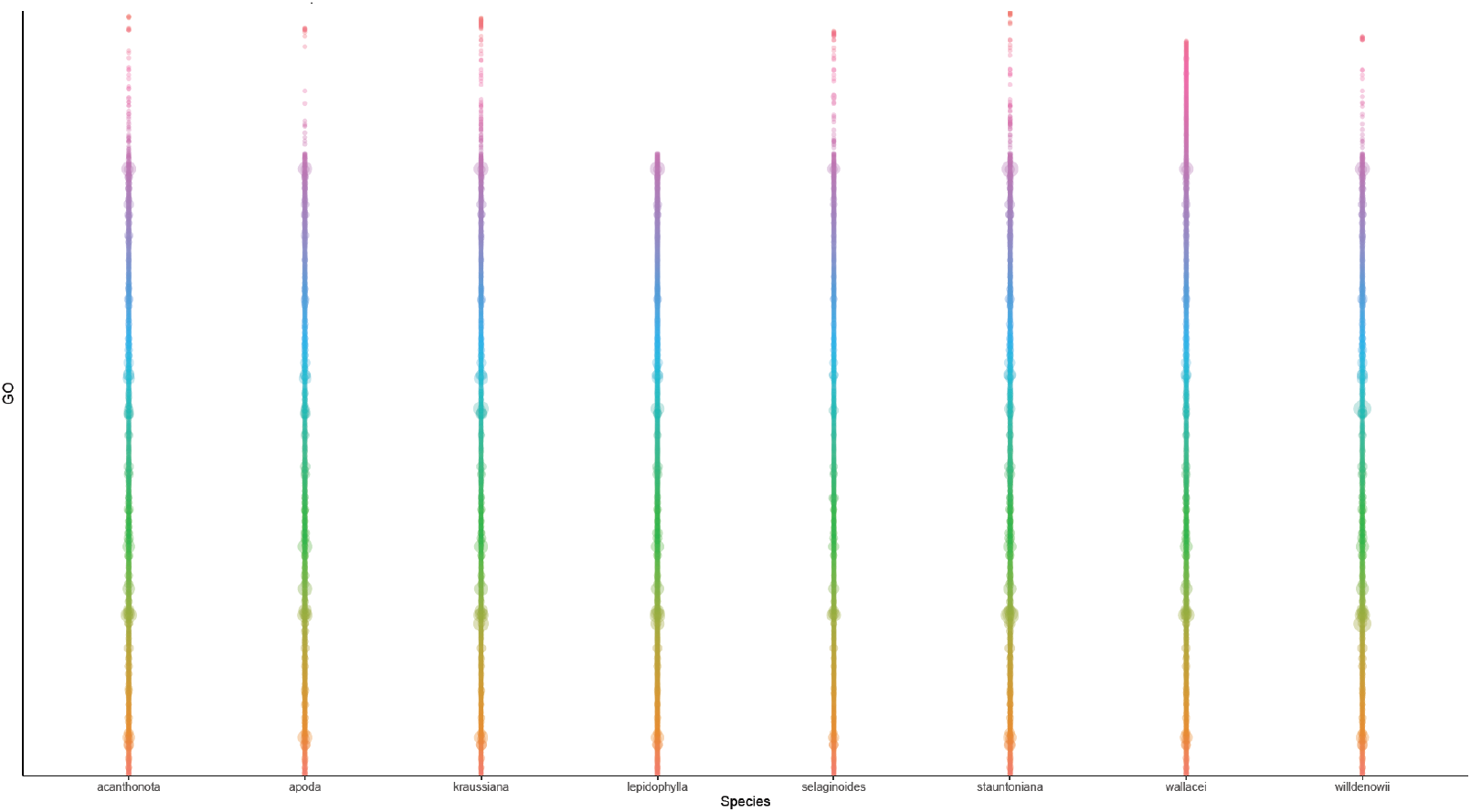
Parallel annotation of GO terms across *Selaginella* species. Visualization created with ggplot2^36^ in R^33,34^.

**Figure 4.**
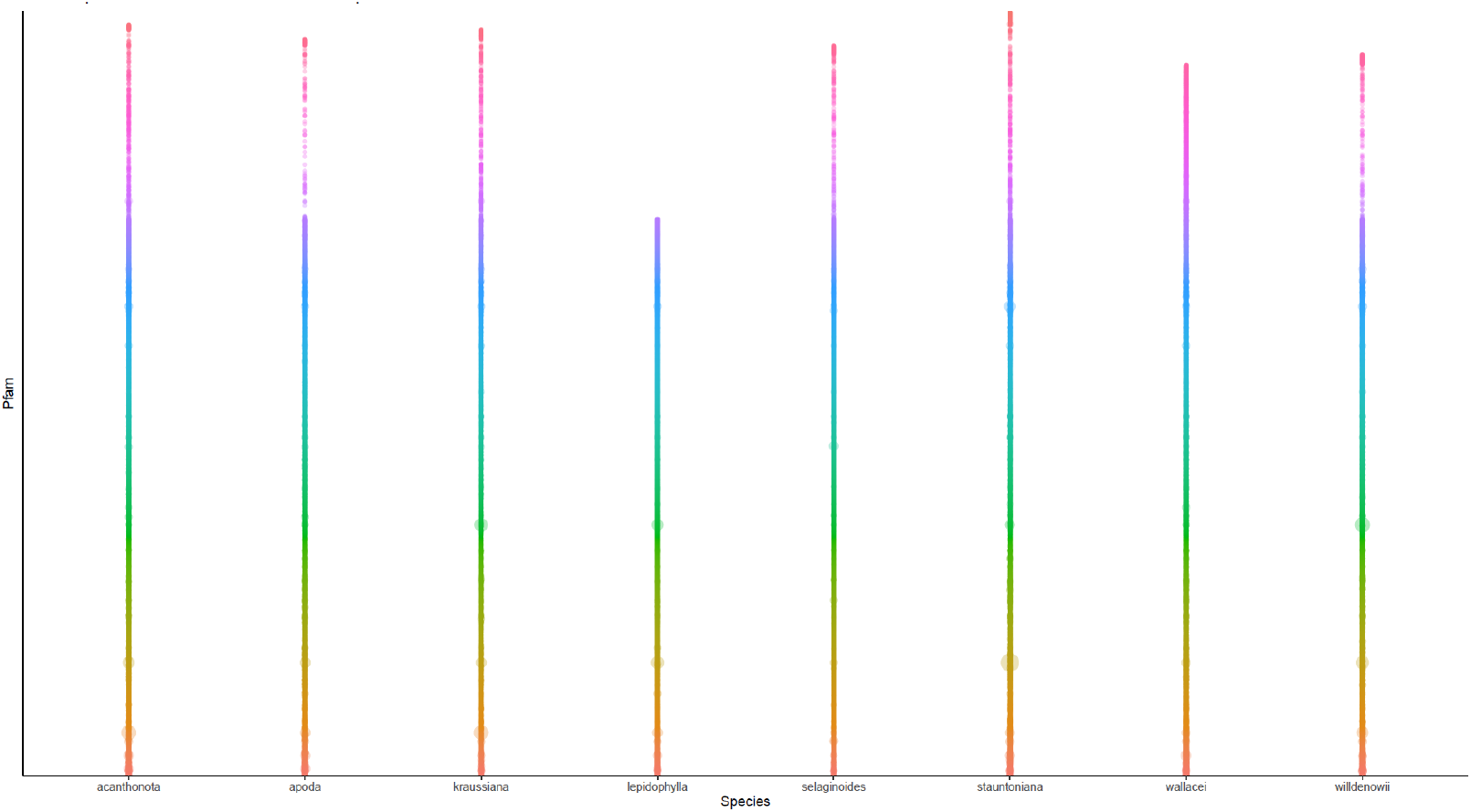
Parallel annotation of predicted Pfam protein domains across *Selaginella* species. Visualization created with ggplot2^36^ in R^33,34^.

We report around 70 to 300 previously unannotated transcripts and predicted proteins. There are proteins with moderate sequence similarity to known *S. moellendorffii* proteins. One discovered protein (PF11421; E-value = 2.1e-4) of ABIJ was found to be >50% identical (E-value = 8e-21) to a mycophenolic acid synthesis protein of *S. moellendorffii*. Another protein (PF01535; E-value = 3.2e-7) of ABIJ was found to be >30% identical (E-value = 9e-147) to a mitochondrial pentatricopeptide repeat protein, a potential RNA editor, of *S. moellendorffii*. Additionally, one protein (PF02224; E-value = 6.8e-11) of ABIJ was >50% identical (E-value = 3e-24) to a cytidylate kinase of *Acinetobacter*, a potential bacterial endophyte, symbiont, or soil contaminant. This protein may have been acquired through horizontal transmission. Alternatively, this protein may have evolved independently. There are at least two predicted proteins that appear to be unlike any other: an antennae protein (PF00556; E-value = 2.5e-4) of KJYC associated with the light-harvesting complex, and a bacterial Abg transporter protein (PF03806; E-value = 1.7e-4) of KUXM potentially associated with folic acid production.

One transcript of KJYC appears to encode a closterovirus coat protein (PF01785; E-value < 1e-4) of *Closteroviridae*, a potential pathogen. The predicted protein shared around 70% sequence similarity (E-value = 3e-102) with a protein found in *S. moellendorffii*. Two predicted proteins of KJYC indicate that root tissue was sampled (PF06830; E-value < 7e-7). The two proteins shared around 70% sequence similarity (E-value = 4e-62 and E-value = 1e-107) with a *S. moellendorffii* protein. One predicted protein (PF08411; E-value = 2.3e-5) of ZYCD was >90% identical (E-value = 6e-40) to a transcription factor associated with a parasitic spider mite, *Tetranychus urticae*. Another protein (PF06330; E-value = 7.9e-60) of ZYCD was >80% identical (E-value = 8e-70) to a trichodiene synthase of *Aspergillus*, a potential fungal pathogen or soil contaminant.

Two predicted proteins of ZZOL had a single-stranded DNA-binding motif (E-value = 1.1e-7 and E-value = 7e-27) associated with telomere protection and were >90% identical (E-value = 3e-49 and E-value < 1e-100) to that of a predicted *S. moellendorffii* protein. The ZZOL sample appears to have significant environmental contamination. One protein (PF15014; E-value = 7e-53) shared >40% sequence similarity (E-value < 1e-40) with that of a fish, anemone, coral, brachiopod, gastropod, bivalve, *Daphnia*, rotifer, worm, aphid, or other insect. Another protein, a pectate lyase (PF03211; E-value = 8.5e-10), shared >70% sequence similarity (E-value < 4e-23) with that of *Arthrobotrys* and *Dactylella*, nematode capturing fungi, or *Xylaria*, a decomposer and potential symbiont. Various other potentially degraded transcripts were also linked to an aquatic environment or recycled aquarium water.

## Discussion

OneKP collaborators identified potential contamination across samples^38^. Initial phylogenetic analyses of *Selaginella* proteins have provided insight on the microbiome. Trehalose is present in various microbes. Trehalose 6-phosphate (T6P) is present in green plants where T6P synthase (TPS) and T6P phosphatase (TPP) catalyze its biosynthesis. *S. lepidophylla* has around the same amount of plant-specific TPP proteins present compared to *S. moellendorffii* and *A. thaliana*. However, *S. lepidophylla* and *S. moellendorffii* have high trehalose content along with high concentrations of microbe-specific TPS proteins, particularly fungi-specific TPS proteins^39^. The microphylls of *S. lepidophylla* contain bacteria and fungi while the xylem have fungal hyphae present among other microbes^11^. And because *S. lepidophylla* is unable to grow from spore when placed in a sterile environment, the authors hypothesized that the microbe-specific proteins are of endophytes^39^. *plant* was able to identify environmental RNA as well as microbiome-specific transcripts. In the future, we can indicate contamination on the visualization. We should also average across replicates to reduce bias associated with haplotype variation.

In essence, *plant* allows for a gene to be compared by contributing its annotations to a pool of annotations. Annotations are universal and can be quantified and compared across all samples. *plant* has many advantages over previous methods, which usually require a sequenced and annotated genome. *plant* is a *de novo* protocol as it requires almost zero prior information about the species. One predicted protein of the model organism *S. moellendorffii* appears to be that of a virus. We propose *plant* as a way to assign taxonomy, classify genes, and clean up the RefSeq nr protein database.

There is one key assumption of *plant*, however. There should be a relatively short evolutionary distance among the species. The rationale is that we want the majority of genes to be expressed similarly in all samples. An alternative is to include a particular amount of a specific RNA transcript in all samples to calibrate the measurements. This consistency ensures that the counts of annotations are normalized properly. If all samples are relatively similar we are then able to identify particularly lowly or highly expressed protein domains via *limma*, which performs an analysis of variance after a TMM normalization across samples^40^. We could then trace the annotations back to the gene-level. Furthermore, we could look at metabolic pathways associated with each gene.

In 2014, Zhu and team refined the mappability aspect of cross-species annotation generation. The authors developed XSAnno to compare neurological activity among well-studied primates with annotated genomes. Their method compared orthologs among species for differential gene expression analysis^41^. This approach essentially compares all “like” components among species. Variation is lost, however. *plant* can potentially capture all aspects.

In 2015, LoVerso and Cui designed a sequential, open-source protocol for cross-species RNA-seq analysis. Their main goal was to select a reference genome and then generate annotated genomes. “Differential expression analysis should focus on those exons and genes that can be measured in all samples.”^42^ Their method is almost *de novo* as it depends on an annotated genome. There are independent studies where collaborators attempt to automate genome annotation^43^. Even then, their comparative method has a focus on the conserved patterns of a gene family and ignores alternative splicing variation among species. *plant*, on the other hand, was built to independently assess each sample and identify unique traits. “Although alternatively splicing is biologically important, comparing all exons in a gene between species is less meaningful as exons may differ in size and number.”^42^ We allow for any exons to be compared indirectly. Each exon contributes its annotations to a pool of total annotations for a sample. *plant* also avoids making very particular biological assumptions about exon usage and splicing.

In 2019, Rey and team developed CAARS (comparative assembly and annotation of RNA-Seq data). CAARS avoids quantifying altogether and constructs gene trees instead. Again, this approach relies on homology among gene families and is unable to isolate species-specific genes. CAARS also depends on assisted assemblies. However, even closely related species are unlikely to map to the same genome properly. The authors note that for “more distantly related species, no approach has been proposed for RNA-Seq assembly”^43^. According to the authors, shared genes become a limiting factor with increasing evolutionary distance. *plant* bypasses the matter by independently assembling each sample *de novo*. With the addition of an RNA spike-in, the abundances of predicted protein domains are always universally comparable.

Other comparative methods focus on shared, core features. We establish a protocol to explore variation. The genes that are left out of these comparisons are actually of particular interest as they form the unique identity of the species. We should have automated pipelines that can fish out new proteins as new ecotypes are sequenced. In fact, greater biodiversity and sample coverage would allow *plant* to disambiguate particular isoforms with greater resolution. Growing the protein database would allow artificial intelligence algorithms to learn at an accelerated rate and help us think of ways to engineer ecosystems to prepare for climate change. With *plant*, we may see the footprints of previously unannotated or undiscovered proteins. The visualization allows us to identify similar patterns of protein domains among species as well as species-specific patterns.

## Future Directions

Next, we could try SOAPdenovo-Trans as the *de novo* assembler for *plant*. With proper parameter tuning, like allowing for an increased number of alternative splice forms compared to just five limited by 1KP collaborators^9^, we could potentially construct a greater number of transcripts. While Trinity takes advantage of dynamic programming, an approach that can find a global optimum, to traverse the Bruijn graphs, Oases^44^ saves time with a heuristic algorithm that has the potential for increased contiguity. SOAPdenovo-Trans attempts to leverage the heuristic approach of Oases with the removal process of Trinity for low redundancy^45^. Another future direction would be to assess RNA secondary structure of the assembled reads. We could then potentially discover novel RNA secondary structure. *plant* is a flexible platform applicable to single-cell RNA-seq data. With *plant*, clustering and prediction algorithms should be able to forecast multipotent stem cell types and differentiate abnormal cells from healthy cells based on similar patterns of protein domain annotations rather than synchronized gene expression. Additionally, machine learning techniques should be able to predict cancer types and prognoses based on annotations rather than genes. If the genomic rearrangements are so great as to render the genes unmappable to the human reference genome, *plant* may have direct applications in medicine.

## Acknowledgments

Asif Ali thanks Barbara Ambrose, Brian Parker, Manpreet Katari, and the HPC team at NYU for support.

## Notes

https://github.com/asifali-bio/plant

### Competing Interest Statement

The authors have declared no competing interest.

## References

1. Keeling, P. J. The endosymbiotic origin, diversification and fate of plastids. Philos. Trans. R. Soc. Lond. B Biol. Sci. 365, 729–748 (2010).

2. Delwiche, C. F. & Cooper, E. D. The evolutionary origin of a terrestrial flora. Curr. Biol. 25, R899–R910 (2015).

3. Tanaka, R. & Tanaka, A. Chlorophyll cycle regulates the construction and destruction of the light-harvesting complexes. Biochim. Biophys. Acta Bioenerg. 1807, 968–976 (2011).

4. The Arabidopsis Genome Initiative. Analysis of the genome sequence of the flowering plant Arabidopsis thaliana. Nature 408, 796–815 (2000).

5. Chen, F. et al. The sequenced angiosperm genomes and genome databases. Front. Plant Sci. 9, 418 (2018).

6. Banks, J. A. et al. The Selaginella genome identifies genetic changes associated with the evolution of vascular plants. Science 332, 960–963 (2011).

7. Gramzow, L. et al. Selaginella genome analysis – entering the “homoplasy heaven” of the MADS world. Front. Plant Sci. 3, 214 (2012).

8. Zhu, Y. et al. Global transcriptome analysis reveals extensive gene remodeling, alternative splicing and differential transcription profiles in non-seed vascular plant Selaginella moellendorffii. BMC Genom. 18, 1042 (2017).

9. Matasci, N. et al. Data access for the 1,000 Plants (1KP) project. GigaScience 3, 17 (2014).

10. Leebens-Mack, J. H. et al. One Thousand Plant Transcriptomes Initiative. One thousand plant transcriptomes and the phylogenomics of green plants. Nature 574, 679–685 (2019).

11. Brighigna, L., Bennici, A., Tani, C. & Tani, G. Structural and ultrastructural characterization of Selaginella lepidophylla, a desiccation-tolerant plant, during the rehydration process. Flora 197, 81–91 (2002).

12. Yobi, A. et al. Metabolomic profiling in Selaginella lepidophylla at various hydration states provides new insights into the mechanistic basis of desiccation tolerance. Mol. Plant 6, 369–385 (2013).

13. Rafsanjani, A., Brulé, V., Western, T. L. & Pasini, D. Hydro-responsive curling of the resurrection plant Selaginella lepidophylla. Sci. Rep. 5, 8064 (2015).

14. Hébant, C. & Lee, D. Ultrastructural Basis and Developmental Control of Blue Iridescence in Selaginella Leaves. Am. J. Bot. 71, 216–219 (1984).

15. Weststrand, S. & Korall, P. Phylogeny of Selaginellaceae: there is value in morphology after all! Am. J. Bot. 103, 2136–2159 (2016).

16. Schulz, C. et al. An overview of the morphology, anatomy, and life cycle of a new model species: the lycophyte Selaginella apoda (L.) Spring. Int. J. Plant Sci. 171, 693–712 (2010).

17. South African National Biodiversity Institute. Selaginella kraussiana. PlantZAfrica http://pza.sanbi.org/selaginella-kraussiana (2013).

18. Plants of the World Online. Kew Science. Selaginella stauntoniana Spring. Royal Botanic Gardens, Kew http://www.plantsoftheworldonline.org (2017).

19. Johnson, M. T. J. et al. Evaluating methods for isolating total RNA and predicting the success of sequencing phylogenetically diverse plant transcriptomes. PLOS ONE 7, e50226 (2012).

20. Leinonen, R., Sugawara, H. & Shumway, M. The International Nucleotide Sequence Database Collaboration. The sequence read archive. Nucleic Acids Res. 39, D19–D21 (2011).

21. Andrews, S. FastQC: a quality control tool for high throughput sequence data. Babraham Bioinformatics http://www.bioinformatics.babraham.ac.uk/projects/fastqc (2019).

22. Grabherr, M. G. et al. Full-length transcriptome assembly from RNA-Seq data without a reference genome. Nat. Biotechnol. 29, 644–652 (2011).

23. Haas, B. J. et al. De novo transcript sequence reconstruction from RNA-seq using the Trinity platform for reference generation and analysis. Nat. Protoc. 8, 1494–1512 (2013).

24. Jones, P. et al. InterProScan 5: genome-scale protein function classification. Bioinformatics 30, 1236–1240 (2014).

25. El-Gebali, S. et al. The Pfam protein families database in 2019. Nucleic Acids Res. 47, D427– D432 (2019).

26. Sonnhammer, E. L. L., Eddy, S. R., Birney, E., Bateman, A. & Durbin, R. Pfam: multiple sequence alignments and HMM-profiles of protein domains. Nucleic Acids Res. 26, 320–322 (1998).

27. Eddy, S. R. Accelerated profile HMM searches. PLOS Comput. Biol. 7, e1002195 (2011).

28. Sonnhammer, E. L. L., Eddy, S. R. & Durbin, R. Pfam: a comprehensive database of protein domain families based on seed alignments. Proteins 28, 405–420 (1997).

29. The UniProt Consortium. UniProt: the universal protein knowledgebase. Nucleic Acids Res. 45, D158–D169 (2017).

30. Finn, R. D. et al. The Pfam protein families database: towards a more sustainable future. Nucleic Acids Res. 44, D279–D285 (2016).

31. Ashburner, M. et al. Gene Ontology Consortium. Gene ontology: tool for the unification of biology. Nat. Genet. 25, 25–29 (2000).

32. Bray, N. L., Pimentel, H., Melsted, P. & Pachter, L. Near-optimal probabilistic RNA-seq quantification. Nat. Biotechnol. 34, 525–527 (2016).

33. R Core Team. R: A language and environment for statistical computing. R Foundation https://www.R-project.org (2019).

34. RStudio Team. RStudio: integrated development for R. RStudio http://www.rstudio.com (2020).

35. Wickham, H. The split-apply-combine strategy for data analysis. J. Stat. Softw. 40, 1–29 (2011).

36. Wickham, H. et al. ggplot2: elegant graphics for data analysis. Tidyverse https://ggplot2.tidyverse.org (2016).

37. Sievert, C. et al. Interactive web-based data visualization with R, plotly, and shiny. Plotly https://plotly-r.com (2020).

38. Carpenter, E. J. et al. Access to RNA-sequencing data from 1,173 plant species: the 1000 Plant transcriptomes initiative (1KP). GigaScience 8, giz126 (2019).

39. Pampurova, S., Verschooten, K., Avonce, N. & Van Dijck, P. Functional screening of a cDNA library from the desiccation-tolerant plant Selaginella lepidophylla in yeast mutants identifies trehalose biosynthesis genes of plant and microbial origin. J. Plant Res. 127, 803–813 (2014).

40. Ritchie, M. E. et al. limma powers differential expression analyses for RNA-sequencing and microarray studies. Nucleic Acids Res. 43, e47 (2015).

41. Zhu, Y., Li, M., Sousa, A. M. M. & Šestan, N. XSAnno: a framework for building ortholog models in cross-species transcriptome comparisons. BMC Genom. 15, 343 (2014).

42. LoVerso, P. R. & Cui, F. A computational pipeline for cross-species analysis of RNA-seq data using R and bioconductor. Bioinform. Biol. Insights 9, 165–174 (2015).

43. Rey, C., Veber, P., Boussau, B. & Sémon, M. CAARS: comparative assembly and annotation of RNA-Seq data. Bioinformatics 35, 2199–2207 (2019).

44. Schulz, M.H., Zerbino, D.R., Vingron, M., and Birney, E. Oases: robust de novo RNA-seq assembly across the dynamic range of expression levels. Bioinformatics 28, 1086–1092 (2012).

45. Xie, Y. et al. SOAPdenovo-Trans: de novo transcriptome assembly with short RNA-seq reads. Bioinformatics 30, 1660–1666 (2014).

46. Soltis, D. The Soltis lab fills the gaps in green plant phylogeny for the Open Tree of Life. Open Tree of Life https://blog.opentreeoflife.org/2013/09/03/filling-the-gaps-in-green-plant-phylogeny-for-the-open-tree-of-life (2013).

47. Fischer, C. Chara. Wikimedia Commons https://commons.wikimedia.org/wiki/File:CharaFragilis.jpg (2005).

